# Preference for Scientist Occupation Among Medical and Science Students in South Korea

**DOI:** 10.1101/2024.01.31.578132

**Authors:** Eun Bit Bae

## Abstract

Although becoming a scientist is generally a long and arduous process, fostering scientists is considered important for national development. To determine medical and science undergraduate students’ preference of scientist, a designed preference survey was administered to 191 medical and 106 science majors, online. The chi-square test, regression, and mediation models were used. The models present significant associations between group and training programs, and between master’s program and scientist preference. Surprisingly, only 24% of the science students were interested in the PhD program compared to the 79.6% of the medical students. Less than 7% in both groups expressed interest in full-time scientist occupation. The study design and findings are newly reported. A negative public consensus of scientist occupation is identified, and master training program shows strong association with choice of scientist occupation. Due to the unstable scientists’ status, a long-term follow-up is recommended afterward program completion for more effectively fostering scientists.

## Introduction

Becoming a scientist or a physician-scientist is generally considered a long and arduous journey full of uncertainty (Ganetzky, 2017; Ikizler et al., 2015; Kopel and Clothier, 2023; Makarewich, 2018; Petersen et al., 2012; Price et al., 2018; Ysseldyk et al., 2019).

Conventionally, scientists have focused on discovering scientific knowledge or developing scientific technologies (Boon, 2020; Jain et al., 2009; Venkatesh and Lipper, 2000; Youtie and Shapira, 2008) to contribute to national economic growth (Anaeto et al., 2016; Borup et al., 2006). Thus, physician-scientist training programs or research programs have been implemented by the government in each nation to foster economic growth, and are often considered important. However, the uncertainty of academic careers and tenured positions for scientists still affects young scientists, and this matter continues to remain a concerning issue (Feldman, 2023; Ganetzky, 2017; Kopel and Clothier, 2023; Martin, 2017; Petersen et al., 2012; Thorp, 2021; Ysseldyk et al., 2019).

Despite the circumstances, there is a great demand for science professionals in various occupations at present because the developments in the convergence science discipline and coming up of new convergence science fields (previously National Academies defined as transdisciplinary integration of life science, physical science, engineering, and etc). The recent rapid growth of convergent technology, such as bio-digital and digital healthcare technology, and its markets are continuing to expand exponentially (Jauffret and Aubrun, 2022; O’Riordan, 2016; Peters et al., 2022). In this current technology development status, the International Electrotechnical Commission (IEC) newly standardized terminology in the discipline of bio-digital convergence (IEC, 2023). Thus, fostering scientists, especially in bio-digital convergence is becoming more crucial, with greater emphasis on implementing programs for their training. Currently, both educational programs for amateurs and training programs for majors and professionals have been internationally implemented in each nation. The U.S. government has successively conducted programs with a long history in interdisciplinary science educational programs, such as biomedical engineering (Linsenmeier and Saterbak, 2020), scientist training programs for summer programs (e.g. Jacksons’ Laboratory since 1924 and the U.S. national program of the Scientific Summer Student Internship Program), and Medical Scientist Training Programs (MSTPs since 1964) from National Institute Health (Harding et al., 2017).

Scientific education and training programs have been conducted in diverse ways led by independent educational institutes and national institutes (Steinman et al., 2020), and entrepreneurs (Aldridge and Audretsch, 2011; Aldridge et al., 2014; Audretsch et al., 2006; Duval-Couetil et al., 2021; Jain et al., 2009; Lehrer and Asakawa, 2004). Most training programs consist of undergraduate programs and national support programs to foster independent scientists in recent convergent science, earlier initiated in biomedical fields (Harding et al., 2017; Linsenmeier and Saterbak, 2020). Through both national/independent programs from institutes and entrepreneurs, training in cutting-edge emerging technologies combined with multidisciplinary techniques are provided to students or scientists.

### Objective and hypothesis

As a part of initiating a project to develop a training program for next-generation scientists, a preference survey was designed to gather statistical data, such as interest in science subjects and inter-semester programs, and willingness to take graduate courses in science, in South Korea. The model for this study was constructed based on the hypothesis that students’ preferences for training programs can affect their occupational preferences as scientists.

## Methods

### Study design

This study design is novel compared to prior studies for the following reasons; 1) the study was implemented on undergraduate students majoring in medicine and various sciences such as biotechnology and engineering, to reflect the current trend of emerging growth in the field of convergence science. 2) the research was implemented before constructing the training program to understand students’ preferences. The preference survey was conducted from August to September 2023, to plan the training project, with the cooperation of four medical colleges, and one another university; the students participated from a total of five universities . The study was conducted using the following procedure. First, I constructed the questionnaire for the preference survey. Second, I screened emerging subjects of convergence science from university’s curriculum for undergraduate students. Third, under the four medical colleges’ cooperation, the preference survey was conducted the medical students via email sent online link; the link was sent to undergraduate students whose majors are medical and science. Fourth, appropriate models were designed based on the characteristics of the data and assessed to examine the significant effects among groups, program preferences, and scientist preferences.

### Construction of the questionnaires

To obtain basal information for developing the training program, the questionnaire for the preference survey mainly included the following six types of questions: 1) interesting subjects in convergence science, 2) inter-semester multidisciplinary programs (i.e., for summer/winter school), 3) master’s degree, 4) doctoral degree, 5) post-doctoral programs, and 6) scientist preference as an occupation. The third to fifth questions are related to research supporting programs involving graduate courses.

The original questionnaires were designed for medical and science students whereby both open-ended and multiple-choice questions consisted of two to ten questions each. From a total of 25 original questions for medical students and 16 questions for science students, essay responses and non-uniform or inconsistent questions were excluded to unify the collected responses into available data. Therefore, categorical data representing the same content from six questions were used in this study.

### Screening convergence science subjects

The post-coronavirus pandemic period, when the planning project was initiated in May 2023, was the fourth year of the COVID-19 pandemic. In this context, digital healthcare and bio-digital technology markets were urgently developed, and convergence science between biotechnology, engineering, and medicine was established as a major in college and undergraduate programs. To develop items for the convergence science question, inter/multidisciplinary programs between biotechnology, engineering, and medicine/public health for undergraduate programs in colleges were investigated, based on which, five convergence scientific subjects were screened (digital healthcare, neurocognitive science, tissue/genetic engineering, big data and AI, and medical devices) and developed as items for the preferred convergence subjects.

### Participants

The participants included undergraduate students recruited from five universities in South Korea (Table S1). For the medical students, four medical colleges of universities cooperated, and for the science students’ group, five science colleges from another single national university were included. The number of science majors was a total of 17, and a detailed list of the science majors is presented in Table S1. To gather participants, a survey notice was posted with an online survey link via an open board at the university, and a message/e-mail was sent with an online link. To promote participation, a mobile gift coupon for a drink was provided within two weeks of completing the survey. A total of 296 undergraduate students (191 medical majors, and 105 science majors) freely participated in this survey voluntarily. The collected data could not be identified personally; age, gender, identification number, and other personal information was not collected (students’ information, which this study can only provide, is grade population; see Table S2). This study corresponds to exemption category 2 in the National Institutes of Health (NIH) determination; thus, formal ethical approval was not required. All methods were performed according to the relevant guidelines and regulations and in accordance with the Declaration of Helsinki and its revised version.

### Statistical analysis

Because the raw data were obtained from two different questionnaires, they were screened mainly for six common types of questions to prepare a model analysis (detailed questions and items in Tables S3 and S4). Driven by six questions, categorical data were coded for statistical analysis. According to the data characteristics, analysis proceeded as follows: first, the Chi-square test was conducted for group comparison and the Cramer’s V was manually calculated and the following formula was obtained (R: number of rows, C: number of columns):

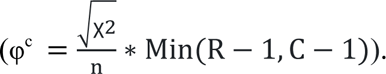

Second, two types of regression models, logistic regression and analysis of variance (ANOVA)-based regression, were evaluated for single-path associations between categorical variables. Third, the mediation model for categorical data was assessed using three variables.

Model construction was based on the main hypothesis that students’ program preference(s) could influence their preference for a scientific occupation. All statistical analyses were performed using SPSS Version 27, IBM, and Process 4.2, which was developed by Andrew F. Hayse and used for mediation analysis.

## Results

A total of 297 participants from five universities (Table S1) and five science colleges from a single institute) were included in the analyses. There were 191 participants from medical colleges and 106 (100 %) from science colleges in the following streams: engineering, 27 (25.5%); biotechnology, 43 (40.6%); information technology, 14 (13.2%); natural science, 13 (12.3%); urban science, 5 (4.7%); and other or unidentified, 4 (3.8%). The differences in the grade populations are presented in Table S2.

### Group comparison results

Descriptive results with frequencies, percentages, and the chi square values of the four parameters are presented in Table 1; Figure 1 shows the frequency of each category of the two groups within a line. Among medical students, the most preferred convergence science subjects were big data and AI (35.6%), brain cognitive science (19.9%), and digital healthcare (13.9%). Science students preferences differed in terms of big data and AI (36.8%), tissue and genetic engineering (22.6%), and brain cognitive science (13.2%). However, the convergence subject preferences between the two groups were not significant (χ2 [6, N = 297]= 8.701, p = .191).

**Figure 1.**
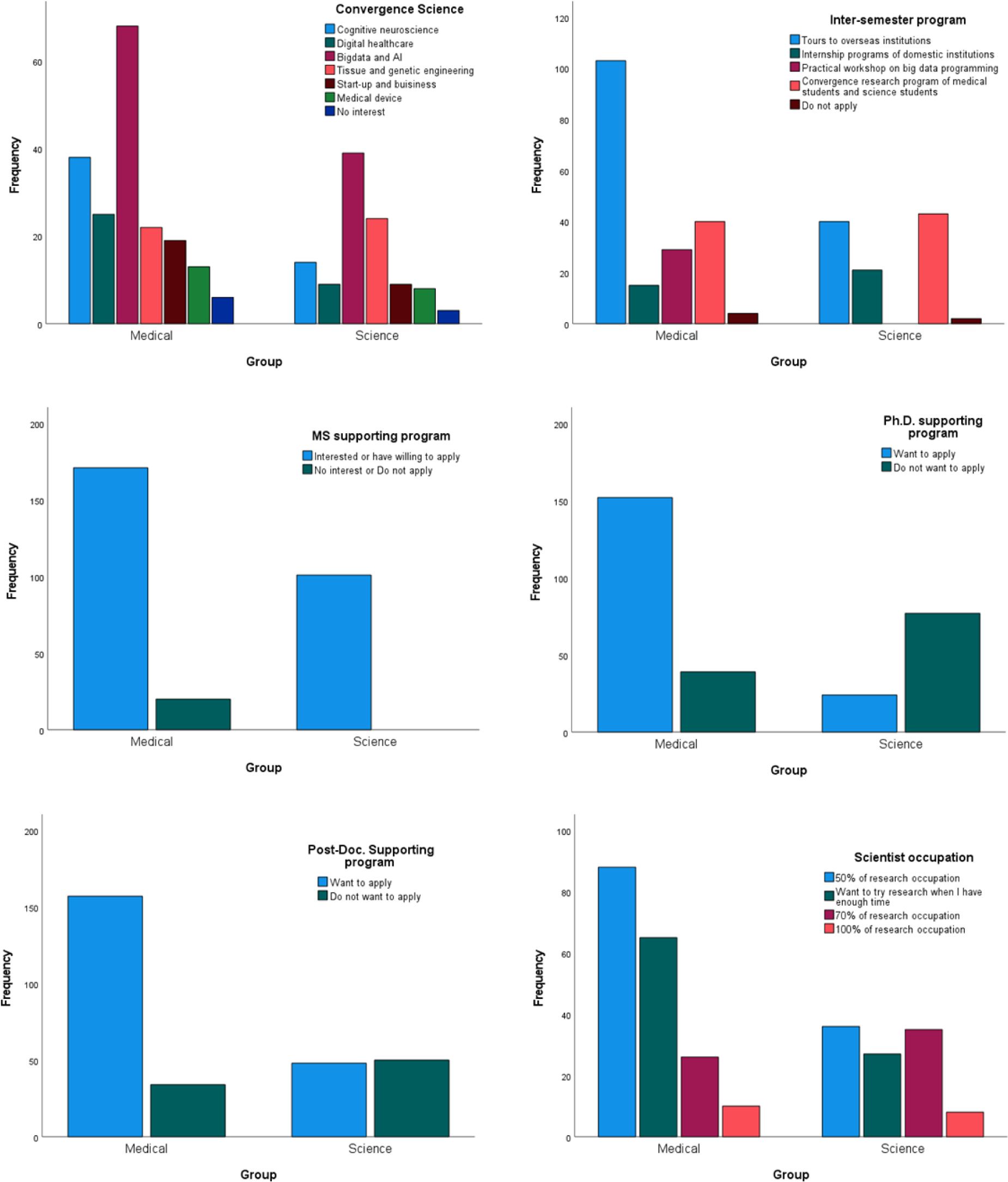
The number of responders is represented in bar graphs classified by group in a line. A total of six subfigures represent responses to six questions.

**Table 1.** The group comparison of the Chi-square test results between medical students and science major students. Among of six questions, five preference questions showed significant differences between the two groups (p <= .001), except convergence science subject.

| **Convergence science subjects** | **Medical** | **Science** | **$\chi^2$ test** |
| --- | --- | --- | --- |
| No interest in science subjects | 6 (3.1) | 3 (2.8) | $\chi^2 (6) = 8.701$<br>$p = .191$<br>$\phi^c = .171$<br>$N = 297$ |
| Digital healthcare | 25 (13.9) | 9 (8.5) |  |
| Cognitive science | 38 (19.9) | 14 (13.2) |  |
| Tissue/genetic engineering | 22 (11.5) | 24 (22.6) |  |
| Bigdata and AI | 68 (35.6) | 39 (36.8) |  |
| Medical device | 13 (6.8) | 8 (7.5) |  |
| Start-up business and management | 19 (9.9) | 9 (8.5) |  |
| Total number of participants | 191 (100) | 106 (100) |  |
| **Inter-semester training program** | **Medical** | **Science** | **$\chi^2$ test** |
| Visits for renowned labs overseas | 103 (53.9) | 40 (41.7) | $\chi^2 (4) = 37.255$<br>$p < .001$<br>$\phi^c = .355$<br>$N = 297$ |
| Domestic internship | 15 (7.9) | 21 (21.9) |  |
| Hands-on workshops on health bigdata programming | 29 (15.2) | 14 (14.6) |  |
| Programs participating both medical and science college students | 40 (20.9) | 19 (19.8) |  |
| Do not apply (No interest) | 4 (2.1) | 2 (2.1) |  |
| Total number of participants | 191 (100) | 96 (100) |  |
| **Scientist training program** | **Medical** | **Science** | **$\chi^2$ test** |
| Interested in master's program | 171 (89.5) | 75 (74.3) | $\chi^2 (1) = 11.354$<br>$p = .001$<br>$\phi^c = .198$<br>$N = 292$ |
| No interest | 20 (10.5) | 26 (25.7) |  |
| Total number of participants | 191 (100) | 101 (100) |  |
| Interested in Ph.D. program | 152 (79.6) | 24 (23.8) | $\chi^2 (1) = 85.966$<br>$p < .001$<br>$\phi^c = .544$<br>$N = 292$ |
| No interest | 39 (20.4) | 77 (76.2) |  |
| Total number of participants | 191 (100) | 101 (100) |  |
| Interested in post-doctoral program | 157 (82.2) | 48 (49.0) | $\chi^2 (1) = 34.666$<br>$p < .001$<br>$\phi^c = .347$<br>$N = 289$ |
| No interest | 34 (17.8) | 50 (51.0) |  |
| Total number of participants | 191 (100) | 98 (100) |  |
| **Scientist occupation** | **Medical** | **Science** | **$\chi^2$ test** |
| a full-time position as a scientist (100% research position) | 10 (5.3) | 7 (6.7) | $\chi^2 (3) = 17.049$<br>$p = .001$<br>$\phi^c = .241$<br>$N = 295$ |
| 70% to research | 26 (13.8) | 35 (33.3) |  |
| 50% to research | 88 (46.6) | 36 (34.3) |  |
| Try to research in my spare time (including 'no interested') | 65 (34.4) | 27 (25.7) |  |
| Total number of participants | 189 (100) | 105 (100) |  |

Visits for overseas field trips were ranked highest in both groups as the most preferred intersemester program. However, there were differences in domestic internship between the two groups; 21% of the science students preferred domestic internships while only 7.9% of medical students preferred this option (χ2 [4, N = 297] = 37.255, p < .001, Table 1).

All the three questions on the research support programs (master’s, doctoral, and postdoctoral programs) showed significant differences between the two groups (p <= .001). Among medical students, 89.5% were interested in the BS/MS combined program, while among science students, 74.3% were interested in the master’s support program (χ2 [1, N = 292] = 11.354, p = .001). Moreover, among medical students, a large percentage of students were interested in the one-year integrated master’s program.

Preference for both doctoral and post-doctoral support programs showed significantly larger differences compared to the master’s program; 79.6% of medical students showed interest in Ph.D. support program while only 23.8% science students were interested in it (χ2 [1, N = 292] = 85.966, p < .001). Also, for post-doctoral support programs, 82.2% medical student and only 29.0% science students showed interest (χ2 [1, N = 289] = 34.666, p < .001, Table 1).

In terms of scientist occupational preferences, the full-time scientist position was ranked the lowest in both groups, in the devotion about 70% to research, only 13.8% medical students responded and the 33% (almost double) of the science students responded. For those who wanted to devote under 70% of their time to research, there were more medical students than science students (χ2 [3, N = 295] = 17.049, p = .001).

### Assessment of Single Path Association

The regression models displayed in Figure 2 were assessed to observe the relationship between the two student groups and program preferences and between program preferences and scientist preference. The results of this model are presented in Table 2. Among the three training programs (master’s, doctoral, and post-doctoral training programs) a single level of logistic regression model showed strong significance in the preference for the Ph.D. training program (χ2 [1, N = 292] = 11.354, p = .000), which explained that medical students were more interested than science students in the Ph.D. training program (β = 12.504, 95% CI = 7.02 - 22.28). For the post-doctoral training program, science students were more interested in 70% and full-time scientist occupations than medical students, while the reverse was true for those who wanted to devote less than 70% of their time to scientist occupations (β = 4.810, 95% CI = 2.80 - 8.28). Based on the above results, among the paths from program preference to scientist preference, only the master’s program model showed significance (F (2, 288) = 4.456, p = .036).

**Figure 2.**
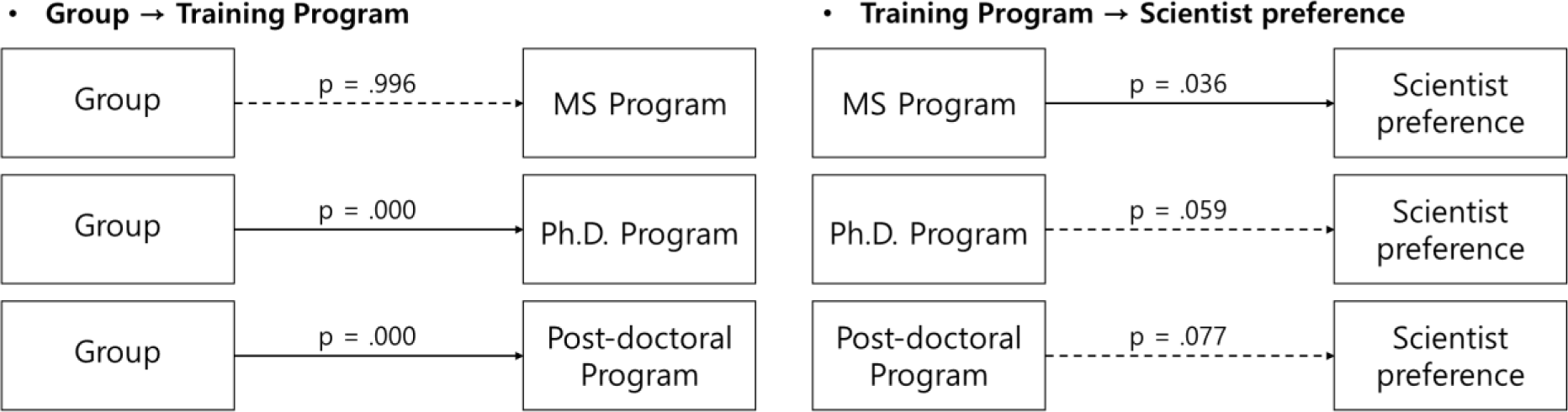
A total of six regression results is represented in the two types of regression models. The single paths from group to training program showed significance in the Ph.D. program and post-doctoral program. The model of training programs to scientist’s preference showed a significant difference in only the master’s program (p = .036).

**Table 2.**
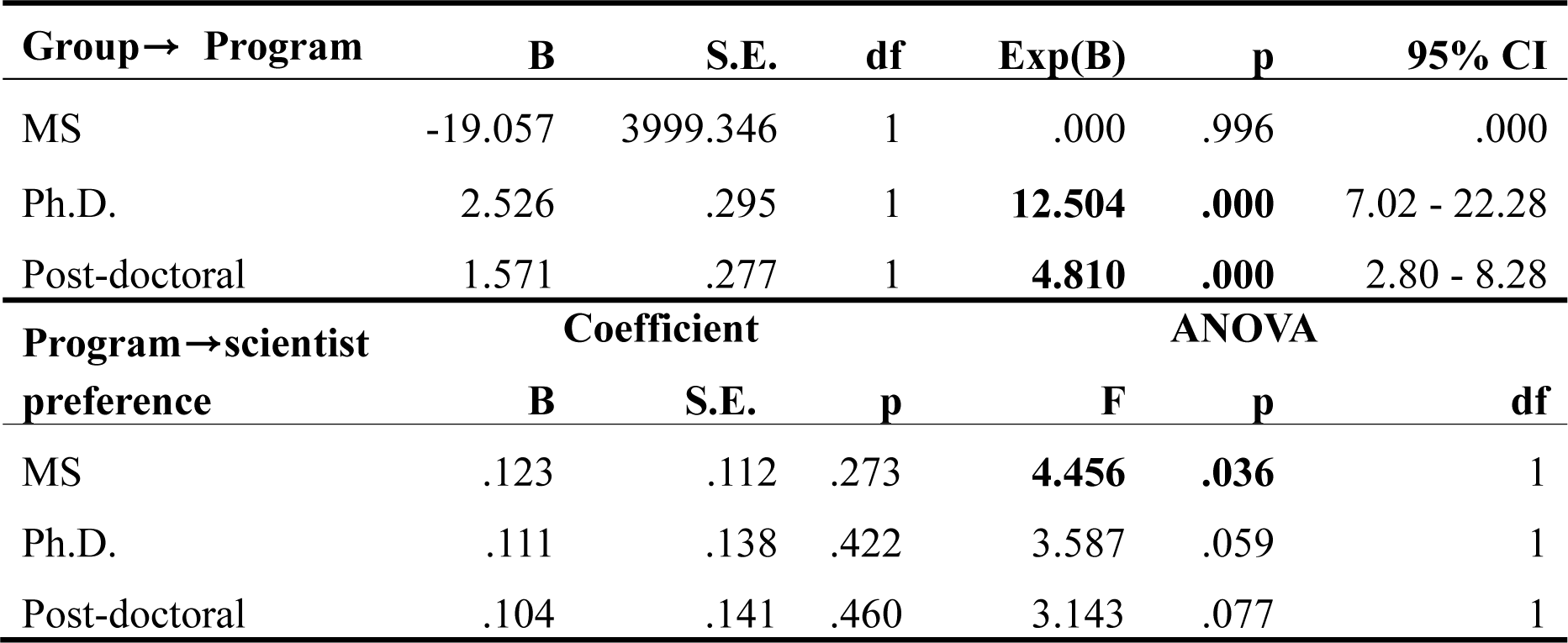
The two types of regression models were analyzed, and six single paths results were presented. The upper model of a single path starts from group to each three of training program preference. Two groups showed differences in both doctoral and post-doctoral training programs. The other model of a single path starts from the training program to the scientist’s preference. Among of three paths, only the master’s program showed a significant difference in the scientist’s preference.

### Assessment of Mediation Model

The associations among group, program, and scientist preferences were assessed in two mediation models: the convergence science subject model and the inter-semester scientific program model. The results are presented in Table 3 and Figure 3.

**Figure 3.**
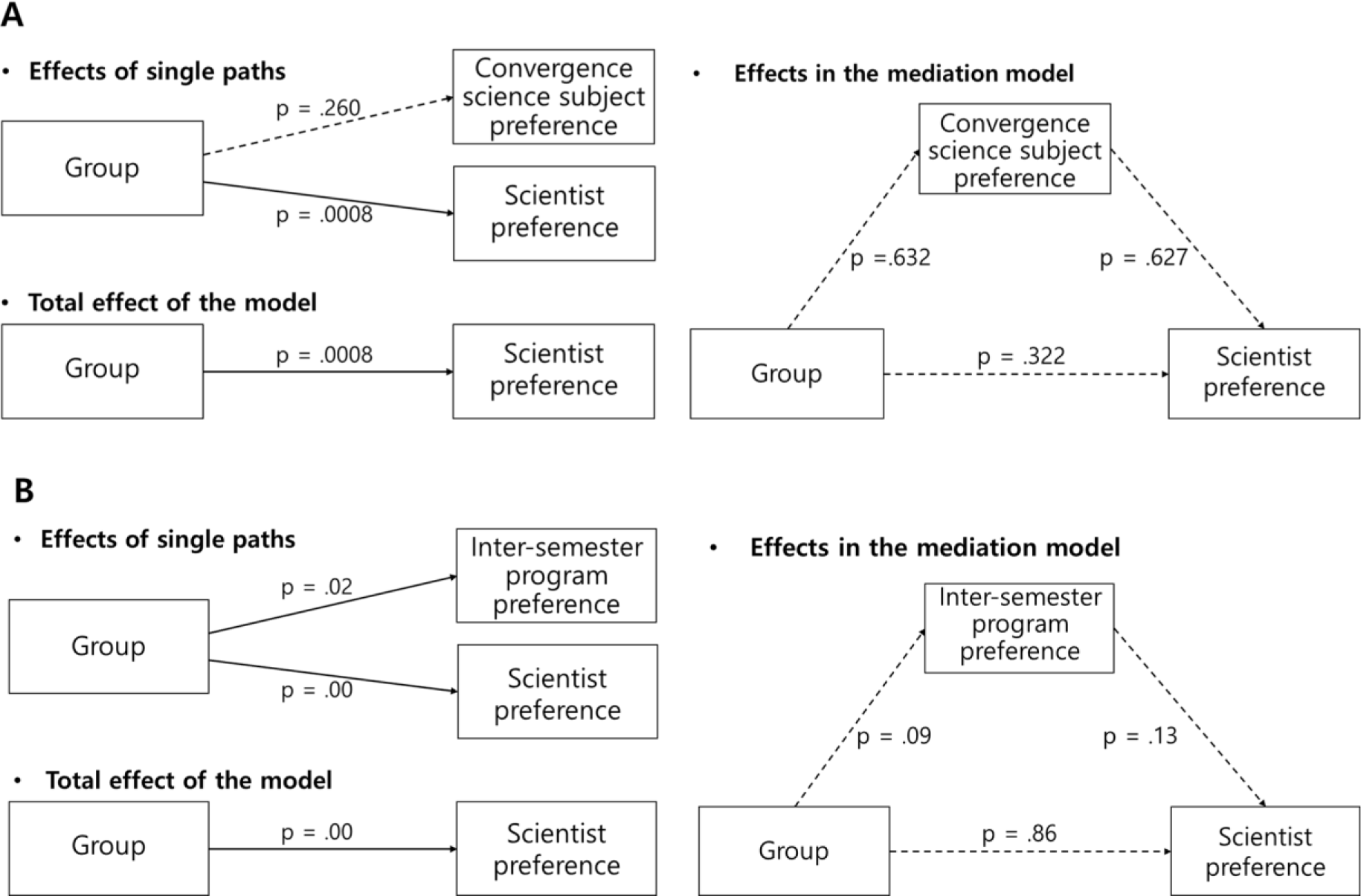
The results of two types of mediation models are represented. While both mediation models were not significant, the total effect from group to scientist preference of both mediation models showed significant results. (p < .001).

**Table 3.** The two types of mediation models were analyzed, and both models showed significant total effects (p < .001). In the upper mediation model, the convergence science subject was a mediator between the group and the scientist’s preference. In the mediation model, the path from group to scientific preference showed significance. In the other mediation model, the inter-semester program was a mediator between group and scientist preference. In this model, two single paths from the group showed significant (p = .02, p = .00).

| **Model for Science subject** | **Coefficient** | **S.E.** | **t** | **p** | **LLCI** | **ULCI** |
| --- | --- | --- | --- | --- | --- | --- |
| **Group → Convergence science subject preference** |  |  |  |  |  |  |
| Constant | 2.896 | .2778 | 10.424 | .000 | 2.349 | 3.443 |
| Group | .220 | .195 | 1.128 | .260 | -.164 | .605 |
| **Group → Scientist preference** |  |  |  |  |  |  |
| Constant | 1.392 | .162 | 8.601 | .000 | 1.074 | 1.711 |
| Group | .386 | .114 | 3.388 | **.0008** | .162 | .609 |
| **Total effect** | **Effect** | **S.E.** | **t** | **p** | **LLCI** | **ULCI** |
|  | .386 | .114 | 3.388 | **.0008** | .162 | .609 |
| **Model for Inter-semester program** | **Coefficient** | **S.E.** | **t** | **p** | **LLCI** | **ULCI** |
| **Group → Inter-semester program** |  |  |  |  |  |  |
| Constant | 1.7 | 0.23 | 7.23 | 0.00 | 1.24 | 2.16 |
| Group | 0.4 | 0.16 | 2.43 | **0.02** | 0.07 | 0.72 |
| **Group → Scientist preference** |  |  |  |  |  |  |
| Constant | 1.41 | 0.16 | 8.83 | 0.00 | 1.1 | 1.73 |
| Group | 0.36 | 0.11 | 3.27 | **0.00** | 0.15 | 0.58 |
| **Total effect** | **Effect** | **S.E.** | **t** | **p** | **LLCI** | **ULCI** |
|  | 0.36 | 0.11 | 2.88 | **0.00** | 0.10 | 0.58 |

In the model for convergence science subject, the direct and indirect effects were not significant (Table S5). However, the single path from group to scientist occupation showed strong significance (Table 3, Figure 3A), and the total effect of the mediation model also showed significance (Effect = .386, SE = .114, 95% CI = .162; .609). The model for the inter- semester program showed no significant direct or indirect effects (Table 3, Figure 3B), while every path showed significant associations between group and inter-semester program preferences (B = .4, SE = .16, t = 2.43, p = .02) and between group and scientist preferences (B = .36, SE = .11, t = 3.27, p = .00). Therefore, the total effect of the mediation model also shows significance from group and inter-semester program preference to scientist preference (Effect = .36, SE = .11, 95% CI = .10; .58).

## Discussion

As presented in the comparison results, a common major interest in convergence science for participants was for big data and AI, which is currently the most in-demand technology across all disciplines. It is considered the most promising and vigorously emerging technology to develop various markets by establishing new converging technologies from diverse fields. Biotechnology ranked second among science students and cognitive science among medical students; these were considered to be affected by external factors such as media, majors, and professors at university. The common interest was overseas visits for field trips which ranked first in both student groups. However, the domestic internship’ program, which ranked second, was considered to significantly affect the total effects of the mediation model between the groups and scientist preferences (p < .001).

The models in this study mainly show newly discovered implications that both majors in the undergraduate period and interest in training programs would affect students career choices as occupations for scientists in undergraduate years. In the regression model between group and scientist training programs, both the Ph.D. and post-doctoral programs were statistically distinguishable between medical and science students (p < .001). Although similar results regarding preference were not recognized in a previous study, Twa et al. (2017) reported that 81% of Canadian MD/PhD graduate physician–scientist trainees were significantly satisfied with the training program (Bianchi et al., 2017). In another study, 81 articles on clinician-scientist training programs were reviewed, and they were found to be dominant in research output compared with intrinsic success (Li et al., 2022). Similar to this study, previous reports have indicated that training programs might have positive effects on scientists’ career choices. To summarize, the model results show that there is a difference in interest in training programs between science and medical students. This significantly affects scientists’ preferences. Another novel finding was that being a scientist or having an occupation in science could be influenced by an undergraduate major or master’s program (Figure 4). In this study, because the total number of participants showed a significant effect of a master’s program on scientist preference, it implies that a master’s training program is more important in determining future careers as scientists or in determining next-generation scientist populations, regardless of college majors.

**Figure 4.**
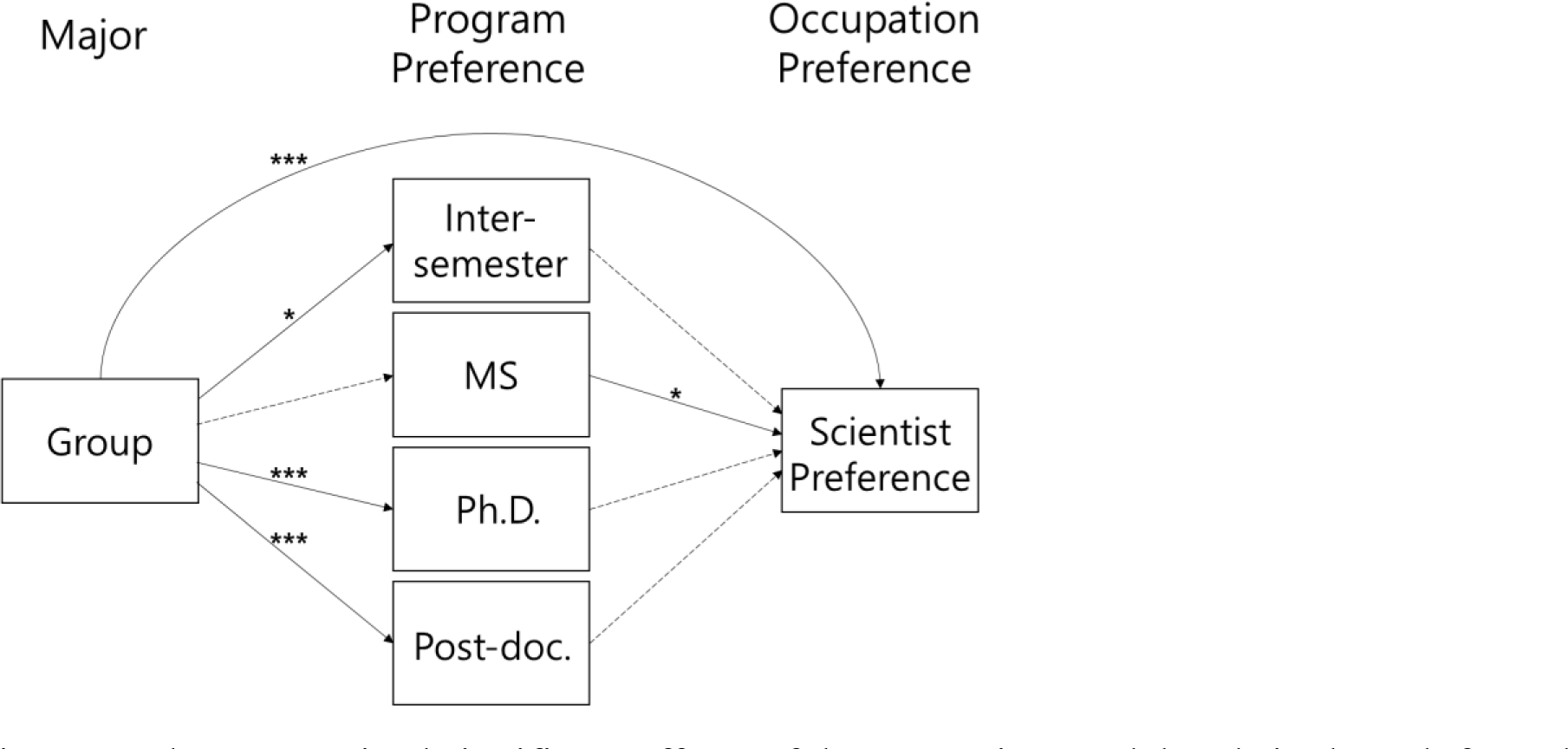
The summarized significant effects of the regression model and single path from the mediation models were represented.

Surprisingly, only 24% of science college students were interested in a Ph.D. programs or in other words, in becoming scientists. Contrarily, 79.6% of the medical students were interested in the Ph.D. program (Table 1). Considering that the preference survey was conducted under the free will of undergraduate students, participants who were interested in being scientists or had favorable attitudes toward being one may have responded to the survey. Under these circumstances, the science group showed paradoxical results of remarkably lower interest in the scientist training program, especially in the doctoral (23.8%), post-doctoral (49.0%), and full-time or part-time scientist occupations. In line with these results, those who preferred a full-time scientist occupation were reported to be less than 7% in both groups, which was the lowest among scientist preferences.

Previously, limitations of scientists’ and young scientists’ careers and positions in academies have often emerged internationally in medical and various science academy associations. The effects of unstable economic and social status of young scientists (Daniels, 2015; Feldman, 2023; Ganetzky, 2017; Ikizler et al., 2015; Lesage and Liston, 2021; Petersen et al., 2012; Price et al., 2018; Schroder et al., 2021; Thorp, 2021) arises once again in the backdrop of this study, with widespread negative public perception that may have spread to university students.

Because the generation of scientists in this era are mainly from fields such as biotechnology, engineering, and medicine, it would not be an exaggeration that future population of scientists among college students also come from these fields. The study was conducted on major’s students before the government of South Korea reported a drastic cut in the upcoming national R&D budget in 2024, which was internationally reported for the first time in 33 years (Jeung, 2023; Normile, 2023; Reardon, 2023). Contrastingly, It was previously reported that the U.S. government requested a 19% increase in the National Science Foundation budget for 2024 to the Congress. Although the preference survey results do not directly reflect the new rising concerns and social emotions regarding the economic and social status of scientists in South Korea, these concerns remain crucial. The continued avoidance of becoming scientists among majors may become a critical limitation in securing competent scientists, and enhancing technological capabilities and competitiveness, directly affecting national economic development.

As previously reported, there is an issue that always comes behind fostering scientists which is unstable economic status and social position for academic scientists. Limited young scientists settling as tenured scientists and instability of their positions in research institutes, academies, and government institutions may interfuse with the public opinion as a social consensus that being a scientist is a long and arduous journey. In sum, the results of this study suggest that a negative public consensus on scientist as an occupation has developed among major’s students. This image can be improved with stable economic and social support and establishing of related regulations by the government.

## Limitations

This study has several limitations. In establishing the questionnaires, several questions and items were commonly constructed; however, the items for the scientist preference question did not seem to fit well, particularly for science college students. For example, a career in science at an academy may be more suitable than that of a full-time scientist. A more plausible language is suggested for further studies according to the characteristics of the group.

Open-ended data were excluded owing to technical and statistical limitations. Therefore, all variables were categorical, which could be a limitation in interpreting the direction of the regression. The data from the training programs were not validated for the mediation models and were then covered under logistic and ANOVA regression models. Another consideration regarding the data is that the science student group was affiliated with a single institute; this could have influenced unknown bias in their responses.

Despite these limitations, there are other expected benefits from this study, such as the statistical results being the base information for estimating the size of the next generation of the scientific and technological workforce, or estimating the application rate when the government promotes various scientist training projects in the future. Considering the limitations of this study, a revised survey in appropriate language is recommended along with a nationally participating study to provide highly reliable results.

## Conclusion

The findings of this study imply that college majors and scientific training programs may influence the preference for a scientist’s occupation. Therefore, countermeasures should be taken to improve the current public perception of the scientist’s occupation in South Korea. After improving the social and economic status of scientist, scientist training programs for next-generation scientists and physician-scientists might be more socially effective.

## Acknowledgements

Thanks to the faculty members of the Seoul National University-Seoul National University Hospital (SNU-SNUH) Physician-Scientist Training Program for their professional advice on the physician-scientist program and the cooperating survey on medical students. Thanks to the science college students at the Incheon National University. Thanks to the Medical Learning Center of the College of Medicine at Korea University, the College of Medicine at Chungbuk National University, and the College of Medicine at Yonsei University for their cooperation in the survey. Thanks to Editage (www.editage.co.kr) for editing and reviewing this manuscript for English language.

